# Rhizosphere bacterial communities of wheat vary across the growing season and among dryland farming systems

**DOI:** 10.1101/700740

**Authors:** Suzanne L. Ishaq, Tim Seipel, Carl J. Yeoman, Fabian D. Menalled

## Abstract

Despite knowledge that seasonality and plant phenology impact soil microbiota, farming system effects on soil microbiota are not often evaluated across the growing season. We assessed the bacterial diversity in wheat rhizosphere soil through the spring and summer of 2016 in winter wheat (*Triticum aestivium* L.) in Montana, USA, from three contrasting farming systems: a chemically-managed no-tillage system, and two USDA-certified organic systems in their fourth year, one including tillage and one where sheep grazing partially offsets tillage frequency. Bacterial richness (range 605 – 1174 OTUs) and evenness (range 0.80 – 0.92) peaked in early June and dropped by late July (range 92 – 1190, 0.62-0.92, respectively), but was not different by farming systems. Organic tilled plots contained more putative nitrogen-fixing bacterial genera than the other two systems. Bacterial community similarities were significantly altered by sampling date, minimum and maximum temperature at sampling, bacterial abundance at date of sampling, total weed richness, and coverage of *Taraxacum officinale*, *Lamium ampleuxicaule,* and *Thlaspi arvense*. This study highlights that weed diversity, season, and farming management system all influence rhizosphere soil microbial communities. Local environmental conditions will strongly affect any practical applications aimed at improving soil diversity and functionality, especially in semi-arid regions where abiotic stress and seasonal variability in temperature and water availability drive primary production.

## 1 Introduction

Microbial communities in agricultural soil are influenced by myriad factors, broadly including seasonality and local environmental conditions, and management practices, and the interaction of these influences is understudied. In a dryland agricultural system, we investigated the interaction of farming system and environmental variables, such as soil moisture and temperature, on the bacterial community associated with the roots of winter wheat (*Triticum aestivum* L.) at various points along a growing season.

Seasonality may result in variations in temperature, precipitation, soil moisture, and solar radiation, all of which drive rates of soil microbial metabolism and respiration, as well as taxonomic composition (Koranda et al., 2013; Orr et al., 2012; Prevost-Boure et al., 2010; Wu et al., 2016). Likewise, environmental conditions alter plant phenology which, in turn, can impact changes in soil microbial communities within a growing season (Donn et al., 2015; Kumar et al., 2018; Sayer et al., 2017). Moreover, management practices, particularly those in agricultural systems, are strongly tied to seasonality (Morrison-Whittle and Goddard, 2015), yet the interplay of time and management on microbial community dynamics in soil has not been well described (Bossio et al., 1998).

Farming practices such as use of pesticides and fertilizers, tillage, crop rotation, irrigation, and the integration of crop and livestock operations directly select for specific microbial communities in soil (Chaudhry et al., 2012; Ishaq, 2017; Lori et al., 2017). Farming practices also indirectly influence soil microbial communities by influencing weed diversity and biomass, soil compaction, soil moisture and permeability, and arthropod populations (Cregger et al., 2012; Lennon et al., 2012; Ponce et al., 2011; Roger-Estrade et al., 2010). Root-associated soil microbial communities can dramatically affect trophic interactions in natural and agricultural settings, by cycling organic material or by interfacing directly with plants to modulate plant growth, root exudates, and health status (Ishaq, 2017; Mariotte et al., 2017). Greater soil microbial phylogenetic diversity is generally considered beneficial for soil and plant health, as it allows for functional redundancy in the provision of ecosystem services, and for the stability and resiliency to disturbances (Bérard et al., 2011; Kuan et al., 2006; Orwin and Wardle, 2004).

Previous studies determined that in comparison with conventional (chemically-managed) no-till and conventional till systems, organic systems have higher soil bacterial cell density and total taxonomic diversity (Chaudhry et al., 2012; Ishaq et al., 2017; Pershina et al., 2015). Yet, many organic systems rely on mechanical tactics like tillage to control weeds. However, tillage use in dryland systems could result in soil erosion, moisture loss, and a change in the community structure of soil microbiota (Ishaq, 2017; Lehnhoff et al., 2017). While there are benefits and limitations of integrated crop-animal production systems (reviewed in (Thiessen Martens and Entz, 2011)), little has been reported on the effect of livestock grazing on soil microbiota. Futher, much of that research has been in pasture-based systems where light stocking increases carbon and nitrogen, and sometimes increases bacterial but not fungal biomass (reviewed in (Ishaq, 2017)).

In the northern portion of the Great Plains of North America, wheat (*Triticum aestivum* L.) is the most widely-planted crop, but production is threatened by insects, climate change, and herbicide-resistant weeds (Keren et al., 2015; Lanning et al., 2010; Menalled et al., 2016). Begininng in 2012, a study has been conducted at the Montana State University Fort Ellis Research and Teaching Center, 3 km east of Bozeman, MT, to compare wheat production challenges across three dryland farming systems: 1) a chemically-managed no-till, 2) an USDA-certified organic system utilizing tillage to manage weeds and terminate cover crops, and 3) an USDA-certified organic system where sheep (*Ovis aries*) grazing is used to manage weeds and terminate cover crops with the goal of reuducing tillage intensity.

In this study, we evaluated patterns in bacterial diversity across the three farming systems described above to elucidate the relative effects of management systems, soil moisture, nutrient content, wheat yield, as well as weed abunandace and diversity on the wheat rhizosphere soil bacterial communities. We hypothesized that 1) date within the growing season be the strongest determinant of bacterial diversity and community structure, 2) farming systems would select for different bacterial communities over the entire growing season, and that 3) farming system would modulate the response of the bacterial community to environmental variables such as low moisture, high temperature, and plant senescence.

## 2 Materials and methods

### 2.1 Site description

Beginning in July 2012, a long-term agricultural field experiment was located at the Montana State University Fort Ellis Research and Teaching Center (45.652664056 N - 110.97249611 W) to assess agronomic and ecological challenges of chemically-managed (i.e. ‘conventional’) and USDA-organic farming systems, as well as the integration of livestock into organic farming systems. Soils at the Fort Ellis site are a Blackmore silt loam (a fine-silty, mixed, superactive, frigid Typic Arguistoll) with 0 to 4% slopes and consistent ratio of 1 part sand, 2 parts silt, 1 part clay by weight (Miller and Menalled, 2015). Monthly air temperature in Bozeman in 2016 was higher than historic maximum and minimums from 1981 – 2010, and mean monthly precipitation (Table S1) was lower by 18 mm in May, 16 mm June, and 14 mm in July (“PRISM Climate Group,” 2018).

The experiment followed a randomized split-plot design; using farming system as the main plot (90 x 75m) with three field replicates per farming system, and crop identity as the split-plot (90 x 13 m). Farming systems consisted of 1) chemical no-till system (CNT), in which synthetic inputs were used in the form of fertilizers, herbicides, and fungicides, 2) USDA-certified till organic (OT), and 3) USDA-certified organic with grazing (OG), which integrates sheep grazing to terminate cover crops and manage weeds, with the overall goal of minimizing tillage intensity in organic production. Split-plots were randomly assigned to a starting crop of a 5 yr crop rotation: year 1 – safflower (*Carthamus tinctorius* L.) under-sown to yellow sweet clover (*Melilotus oficinalis* (L.) Lam.), year 2 – sweet clover cover crop, year 3 – winter wheat (*Triticum aestivum* L.), year 4 – lentil (*Lens culinaris* Medik.), and year 5 – winter wheat (Lehnhoff et al., 2017).

Chemical inputs utilized in the CNT system included 2,4-D, bromoxynil, dicamba, fluroxypyr, glyphosate, MCPA, pinoxaden, and urea for winter wheat rotations [see Tables 2.7 and 2.8 in (Johnson, 2015)], which are reflective of typical farm management practices in the Northern Great Plains region. Both organic treatments began the organic transition process in July 2012, making crops harvested as of 2015 USDA-certified as organic. In the OT system, tillage was accomplished using a chisel plow, tandem disk, or field cultivator, as needed for weed control, seedbed preparation, and to incorporate cover crops and crop residues. Weed control was enhanced with a rotary harrow. In the OG system, targeted sheep grazing was used to reduce tillage intensity for pre-seeding and post-harvest weed control and to terminate the cover crops, with duration and intensity of grazing based on weed biomass (Lehnhoff et al., 2017). Grazing was supplemented with tillage as necessary, based on soil conditions and weed pressure. Seeding was done with a low-disturbance no-till double-disk seeder. Further details of the management practices used within each system and farming history prior to planting of these crops can be found elsewhere (Barroso et al., 2015; Johnson, 2015; Lehnhoff et al., 2017). Outside of normal farm management activities, soil disturbance and compaction was minimized during sampling procedures.

### 2.2 Soil measurements and collection

In the present study, soil was sampled in the year 3 winter wheat split plots of each faming systems. Soil moisture was measured weekly using gypsum blocks (Friis Dela, 2001) buried in the center of each sample area to a depth of 7.5 cm and a Delmhorst soil moisture tester (Model KS-D1, Delmhorst Instrument Co.). Every four hours between April 14, 2016 (one week prior to the first sampling) and July 25, 2016 (final sampling date), soil temperature was recorded within sample areas using an iButton (Maxim Integrated) buried at 7.5 cm.

To characterize soil microbial communities, rhizosphere soil cores were obtained from each one of the nine year 3 winter wheat split-plots, within an 0.75 m^2^ area, situated randomly along the length of each 90-m split-plot. Three soil cores were obtained to a depth of 15 cm using a 2-cm diameter core sampler, which was sterilized with 70% isopropanol and air-dried between sample areas. and from each sample area. Soil (50 mL) was homogenized and placed on ice until transport back to Montana State University, where they were stored at −20°C until analysis. Each area was repeatedly sampled five times during the 2016 growing season: April 21, May 12, June 1, June 22, and July 25 (prior to wheat harvesting). Additional soil samples were obtained on July 25, and stored at 4°C until shipped to an independent laboratory (Agvise Laboratories, Northwood, North Dakota, US) for analysis of organic matter, nitrate, phosphorous (Olsen), potassium, and pH (Table S2).

### 2.3 Plant community measurements and collection

Aboveground biomass of all weed species present within sampled areas was harvested by hand in late June, when most weeds and wheat had matured. Weeds were visually identified and separated by species, dried in an oven at 55° C for two weeks, and weighed. Total wheat biomass was harvested from sampled areas by hand from 1.5 row meters on July 25, 2016, immediately after soil samples had been collected. Wheat biomass was dried in an oven for a week at 55° C, weighed, and then mechanically threshed to remove grain. The grain was submitted for protein analysis to the Montana State University Grain Quality Lab (Bozeman, MT).

### 2.4 DNA extraction and sequencing

DNA was extracted using 0.25 g of a 15 - 30 g soil sample homogenized from at least 3 soil cores within the sample area and processed following protocols described in Ishaq et al. (2017) using the PowerSoil 96-well Soil DNA Isolation Kit (MoBio Laboratories, Inc.). Following extraction, an additional cleaning step was added: a 10% volume of 2M sodium acetate was added to each sample, followed by a 200% volume of 100% ethanol. Samples were vortexed and refrigerated overnight at −20°C to precipitate DNA, after which they were centrifuged at 16,000 x G for 5 min, supernatant was poured off, and sample tubes were air-dried. Pellets were washed with 100% ethanol, allowed to air dry again, and eluted into 100 μl of molecular-grade water. The V3-V4 region of the 16S rRNA gene was PCR amplified using the KAPA HotStart PCR Kit (Kapa Biosystems, Wilmington, MA) with 10 µL Kappa HotStart Mastermix, 6 µL molecular-grade water, 1 µL of each forward and reverse primer at 10 mM concentration, and 2 µL sample DNA. PCR protocol was as follows: 95° C for 3 min; 5 cycles of denaturation at 98° C for 20 sec, annealing at 52° C for 30 sec, elongation at 72° C for 45 sec; 25 cycles of denaturation at 98° C for 20 sec, annealing at 60° C for 30 sec, elongation at 72° C for 45 sec. Primers included the MiSeq adaptors (A for forward, B for reverse), the sample index/barcodes, the two-nucleotide linker, and primers 341F (5’-CCTACGGGAGGCAGCAG-3’) and 806R (5’-GGACTACHVGGGTWTCTAAT-3’) (Caporaso et al., 2011). High-throughput sequencing was performed using an Illumina MiSeq (Illumina, San Diego, CA) and a 500-cycle V2 kit, with PhiX used as a positive control at a 10% spike-in, and molecular-grade sterilized water as a negative control. Sequencing output data can be found in the Sequence Read Archive (SRA) at NCBI under BioProject PRJNA383161.

### 2.5 DNA data processing and analysis

Bioinformatics were performed similar to previously described protocols (Ishaq et al., 2017) with a few exceptions. Forward and reverse DNA sequence fragments were assembled into contigs using PANDAseq (Masella et al., 2012) with > 15 nucleotide overlap and default quality parameters, then processed using mothur ver. 1.38 (Schloss et al., 2009). Sequences that contained ambiguous bases, homopolymers > 8 nt, < 300 nt or > 580 nt, were discarded. Sequences were aligned to the Silva nr 119 database (Quast et al., 2013) using Needleman-Wunsch alignment (Needleman and Wunsch, 1970), after which all-blank columns and unaligned sequences were removed. Samples with > 50,000 sequences were subsampled down to 50,000 to improve analysis time. Chimeras were identified and removed using mother-integrated UCHIME (Edgar et al., 2011), and then taxonomically classified with the Wang algorithm/Ribosomal Database Project (RDP) Classifier (Wang et al., 2007) and Silva reference database. Sequences identified as unknown, mitochondria, chloroplast, or Eukaryotic (< 500 combined) were removed. Sequences which were identified to genera known to contain nitrogen-fixing species in soil were identified from previous literature. The number of seqences per sample which passed QA ranged from 1,305 – 157,761.

Sequences were subsampled (normalized) to the size of the smallest sample (n = 3232 sequences per sample), removing nine samples with fewer sequences each from downstream analysis. Distance was calculated considering consecutive gaps to be one event, and sequences were clustered into presumptive species-level operational taxonomic units (OTUs) using the nearest neighbor algorithm at a 0.03 cutoff. OTUs which only contained singletons or doubletons were removed. One sample sampled in July in OT (OT_725_114) contained a dramatically different microbial community than its replicates as it had more Firmicutes and Bacteroidetes and far fewer of any other phyla represented than other samples (included in Fig 4, S2), and was removed from all group-based statistical analysis.

Diversity was assessed using the mothur-integrated versions of Shannon Diversity (Segata et al., 2011), and multivariate analysis and data visualization was performed in R (RCoreTeam, 2018). Diversity data did not meet normality distribution assumptions via Shapiro-Wilks test, thus comparisons of means were conducted with non-parametric Conover tests (Dinno, 2017) using Bonferrroni p value correction for multiple comparisons. Classification-based random forest trees with permuational analysis (rfPermute) were used to identify discriminant taxa, based on “mean decrease accuracy” or the importance of a factor in predicting the microbial community, as measured by randomly re-assigning factor levels and comparing to the original tree of microbial community distribution. Regression-based random forest trees were used to identify important factors for Shannon diversity of abundance of specific taxa (Breiman et al., 2018). The number of trees which minimized the out-of-box error rate was used, typically ntree = 500.

Unweighted Jaccard distance (uJ) and Bray-Curtis Dissimilarity (BC) were calculated to compare community membership (presence/absence) and community structure (presence/absence and abundance), respectively, and assessed using permutational analysis of variance (PERMANOVA) with the Adonis function in the vegan package (Oksanen et al., 2012), or pairwise in the RVAideMemoire package (Herve, 2019), with Block to stratify data, split-plot as a repeated measure, 1,000 permuations, and Bonferroni p-value correction. There was heterogeneity in samples by replicate block, indicating a spatial difference in the soil conditions, which significantly affected bacterial communities (PERMANOVA, uJ, F = 4.498, *p* = 0.001; Bray-Curtis, F = 3.6676, *p* = 0.001), thus, all permutational calculations of distance included field replicate block as a stratification. Non-Metric Multidimensional Scaling Plots (NMDS) based off Bray-Curtis Dissimilarity were calculated in mothur and were visualized using ggplot2 (Wickham, 2009). A heatmap of significant Pearson’s correlations between treatment parameters and OTU abundance was created using corrplot (Wei et al., 2017), which generated correlations and tested significance.

Distance-based redundancy analysis (dbRDA) was conducted using Helligner-transformed community data (Legendre and Gallagher, 2001) and the capscale function of the vegan package, using Block as a condition. The environmental data included categorical treatment levels; soil moisture at soil sampling; the minimum, maximum, and mean soil temperature on the day of soil sampling; the minimum, maximum and mean soil temperature averaged over 3 and 7 days prior to sampling. Biotic and abiotic (meta-) data also included bacterial Shannon diversity; total weed species, total weed coverage, individual weed species coverage, and wheat coverage the previous fall (October 25, 2015), early spring (April 8, 2016), and mid-summer (June 14, 2016), as well as weed biomass in mid-summer (June 14, 2016) and wheat biomass in late-summer (July 25, 2016). Additional significant variables were removed from the original model to create a reduced model, as they were co-linear aliases of included variables, including the biomass in June 2016 of *C. pastoris*, *C. arvense*, *Melilotus* spp., and *T. arvense*, the plot coverage in April 2016 of *C. album*, *G. aparine*, *L. serriola*, and *T. arvense* plot coverage the previous fall in October 2015.

Each farming system contained a distinct weed community profile, and collectively the dominant weed species identified over all systems and time points included *Bromus tectorum* L.*, Capsella bursa-pastoris* L.*, Chenopodium album* L.*, Cirsium arvense* L.*, Galium aparine* L.*, Lactuca serriola* L.*, Lamium amplexicaule* L.*, Lens culinaris* (Medikus)*, Malva neglecta* (Wallr.)*, Melilotus officinalis* L.*, Monolepsis nuttalliana* (Schult.)*, Taraxacum officinale* L.*, Thlaspi arvense* L.*, Tragopogon dubius* (Scop.)*, Trifolium hybridum* L., and *Trifolium pratense* L. Factors for which only one time point was present, such as wheat biomass, soil pH, and soil nutrients, all collected at harvest, were assessed for the July time point only. Additional information and analysis on wheat and weed data are provided elsewhere (Lehnhoff et al., 2017; Seipel et al., 2018). Across all tests, significance was determined at *P <* 0.05

## 3 Results

The bacterial community in soil was comprised of Actinobacteria, Proteobacteria, and Acidobacteria, followed by Bacteroidetes, Firmicutes, Gemmatimonadates, Verrucomicrobia, and Planctomycetes, and several low-relative-abundance phyla (Fig. 1). Within each sampling date, the relative abundance of Acidobacteria, Actinobacteria, Armatimonadetes, Chlorobi, Chloroflexi, Cynaobacteria, Deinococcus-Thermus, Fibrobacteres, Firmicutes, Proteobacteria, and Verrucomicrobia, as well as the candidate phyla BD1-5, BRC1, SHA-109, SM2F11, TM7, WCHB1-60, and WS6 differed among farming systems (Fig. 1; Table S3, *p <* 0.05). There was an increase in the mean coefficient of variation for the relative abundance of each phylum over the growing season (Fig. S1), indicating that there was more variation in field replicates as bacterial abundance was divergent across individual sample areas when soil was hottest and driest in July.

**Figure 1.**
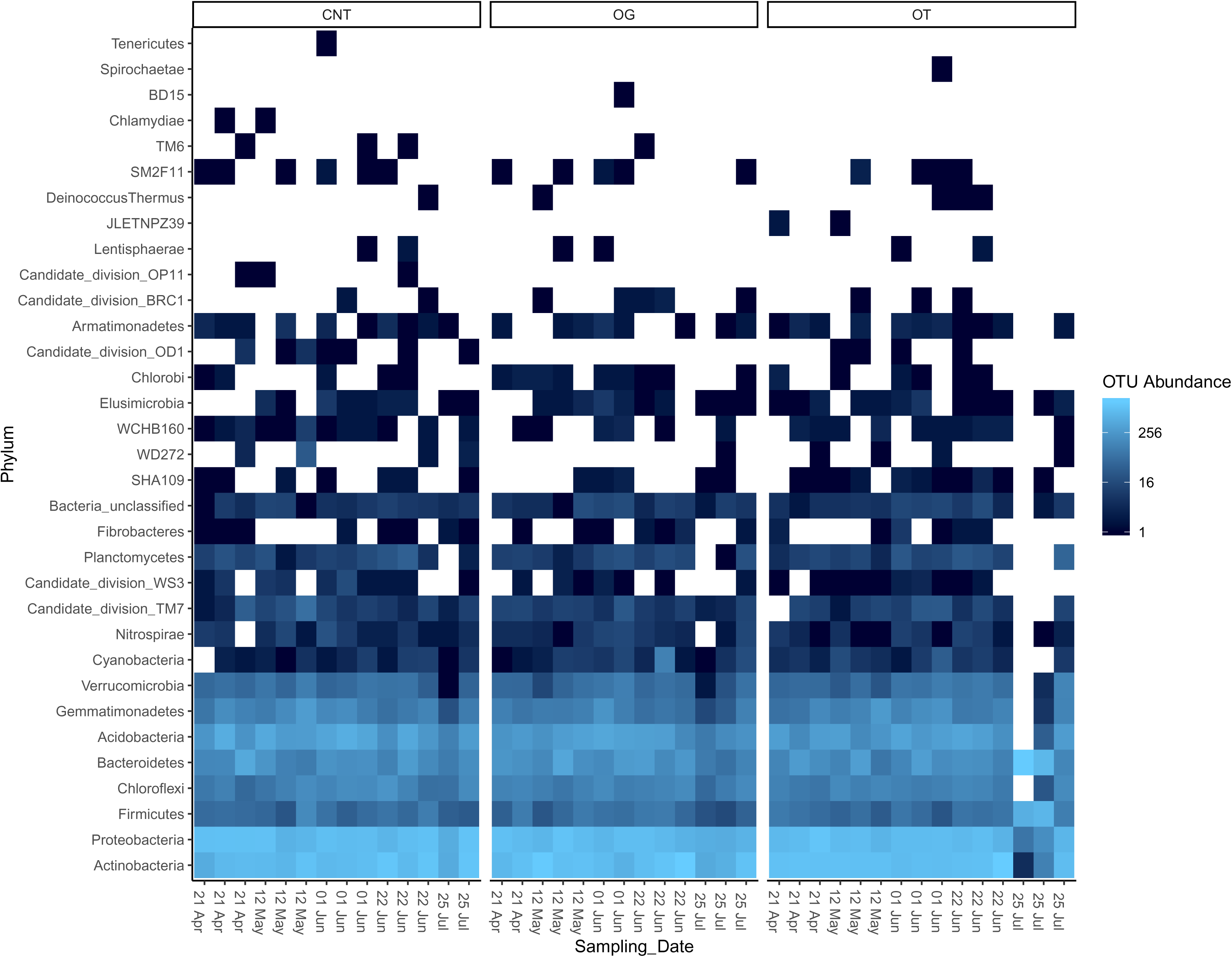
Relative abundance of bacterial phyla in soil from conventional (CNT), organic grazed (OG), and organic tilled (OT) systems during the 2016 growing season. Samples are grouped along the x-axis by farming system, and sorted by sampling date within the growing season.

A total of 8,547 OTUs (97% cutoff) were identified, including many common soil genera which were abundant throughout the growing season in all plots, including *Blastococcus*, *Arthrobacter*, *Skermanella*, *Sphingomonas*, as well as unclassified genera from several different families (Fig. S2). Other OTUs were specific to system, including *Bacteroides* which was prominent in OT sample 202 during July, and the insect-associated *Wolbachia* which was abundant only in a single CNT sampled area in May (Fig. S2).

### 3.1 Seasonal effect

The effect of sampling date was manifested by the unimodal trend of soil bacterial richness over the 2016 growing season. Across the three farming systems, observed OTU (97% cutoff) richness peaked in early June, when it was greater than in April, May, or July (Fig. 2 A, B; Conover, *p* < 0.05 each, Bonferroni corrected (BF)). Richness was also greater in late June than in July (Fig. 2 A, B; Conover, *p* < 0.05 each, BF). Similarly, Shannon-Weiner diversity index of soil bacteria peaked in early June when it was significantly higher than in April, May, late June, and July (Fig. 2C; Conover, *p* < 0.05 each, BF). This change in Shannon diversity over the growing season was driven by changing richness, rather than changes in evenness (Fig. 2D) which was only significantly increased in early June as compared to April (Conover, *p* = 0.004, BF).

**Figure 2.**
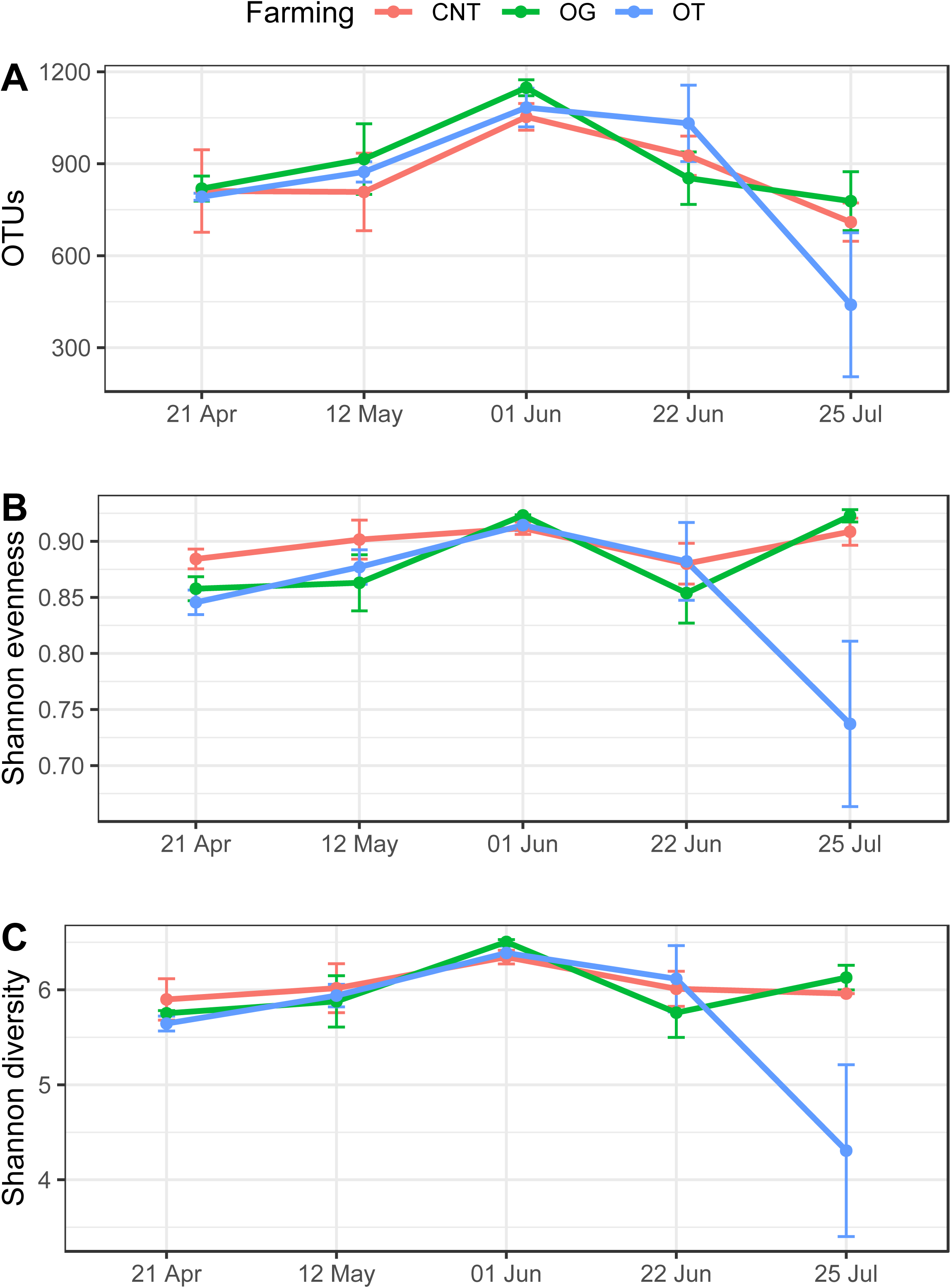
Soil bacterial communities’ A) richness, B) evenness and C) diversity indices for from conventional (CNT, red), organic tilled (OT, blue) systems, and organic grazed (OG, green). Error bars show Standard Error of Means (SEM) for n = 3 samples per time point and farming system.

Sampling date significantly affected bacterial community similarity based on member presence/absence collectively across the growing season (Table 1, uJ), but was not significantly altered in any pairwise sampling date comparison (uJ, *p* > 0.05 BF), contrary to our first hypothesis. Sampling date similarly affected bacterial community similarity in relation to relative abundance (Fig. 3; Table 1, BC, *p* = 0.0001), and was driven by significant differences between June and July (Table S4) when richness was low: early June - July (*p* = 0.014), and late June-July (*p* = 0.022).

**Figure 3.**
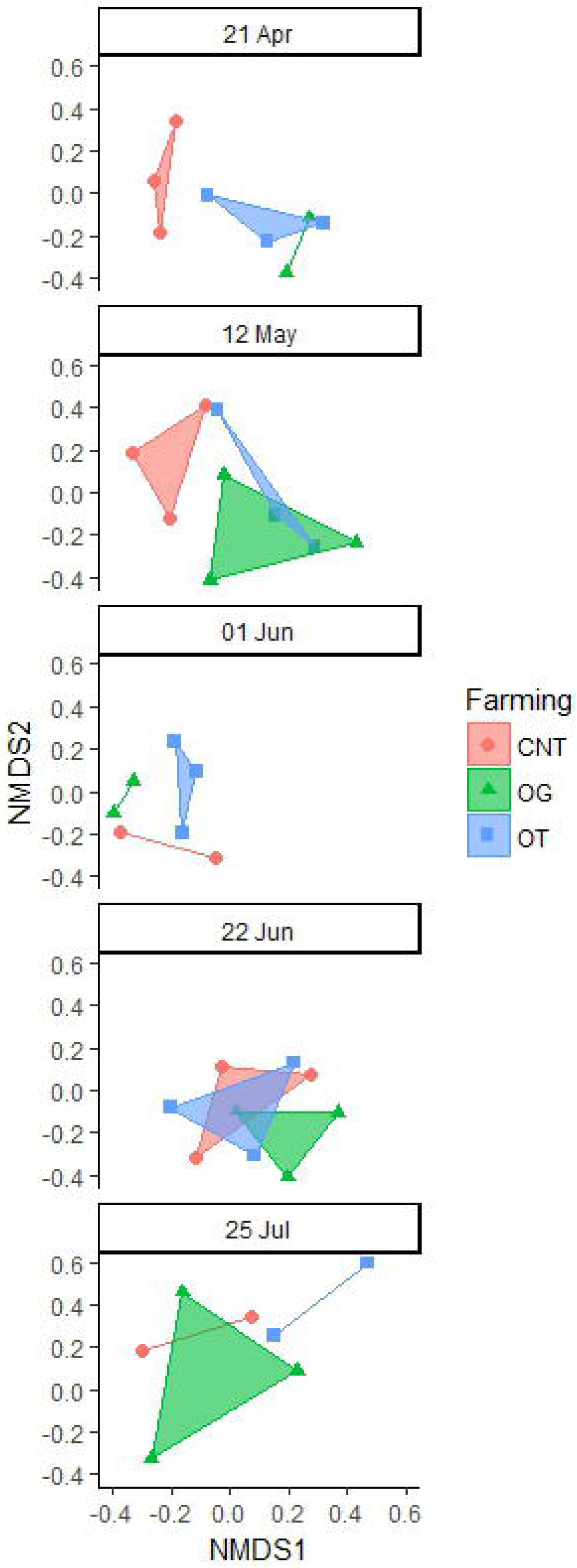
Non-Metric Multidimensional Scaling (NMDS) of Bray-Curtis dissimilarity for soil bacterial communities from conventional (CNT), organic tilled (OT), and organic grazed (OG) systems, over the 2016 growing season.

**Table 1.** PERMANOVA model output of the effects of farming systems, date of sampling and their interactions on soil microbial communities for OTUs at a 97% genetic cutoff.

| Jaccard Simmilarity (unweighted) |  |  |  |  |  |  |  |
| --- | --- | --- | --- | --- | --- | --- | --- |
|  | Df | SumsOfSqs | MeanSqs | F.Model | R2 | Pr(>F) |  |
| Farming | 2 | 0.8421 | 0.42103 | 1.36298 | 0.06524 | 0.0001 | *** |
| Date | 4 | 1.4228 | 0.3557 | 1.15148 | 0.11024 | 0.0009 | *** |
| Farming:Date | 8 | 2.3009 | 0.28761 | 0.93107 | 0.17828 | 0.5954 |  |
| Residuals | 27 | 8.3404 | 0.3089 |  | 0.64624 |  |  |
| Total | 41 | 12.9061 |  |  | 1 |  |  |
| Bray-Curtis Dissimilarity (weighted) |  |  |  |  |  |  |  |
|  | Df | SumsOfSqs | MeanSqs | F.Model | R2 | Pr(>F) |  |
| Farming | 2 | 0.6692 | 0.33458 | 1.66846 | 0.07503 | 0.0002 | ** |
| Date | 4 | 1.2354 | 0.30886 | 1.54019 | 0.13852 | 0.0001 | ** |
| Farming:Date | 8 | 1.5995 | 0.19994 | 0.99705 | 0.17935 | 0.477 |  |
| Residuals | 27 | 5.4144 | 0.20053 |  | 0.60709 |  |  |
| Total | 41 | 8.9185 |  |  | 1 |  |  |

### 3.2 Farming system

Farming system had an equivocal effect on community membership to sampling date during the growing season, and significantly affected both OTU presence/absence (Table 1, uJ), and membership with respect to relative abundance (Table 1, BC). The collective effects that farming system exerted upon soil bacterial communities was observed between CNT and OG plots (Table S5), both in presence/absence (uJ, F = 1.387, *p* = 0.01), and weighted community structure (BC, F = 1.676, *p* = 0.014), indicating a difference in presence/absence as well as relative abundance, respectively. OG and OT contained significantly different communities by presence/absence (uJ, F = 1.41, *p* = 0.011). CNT and OT had only equivicoally different presence/absence composition (uJ, *p* = 0.051).

Richness, evenness, and Shannon Diversity were not significantly different between farming systems at any time point, or averaged across the growing season (Conover, *p* > 0.05, Bonferroni corrected (BF)). Thus, there was no significant difference on a pairwise basis between each farming system within each time point (Fig. 3; PERMANOVA uJ and BC, *p* > 0.05). However, there was more variability in soil bacterial communities at the end of the growing season (betadisp, uJ and BC, *p* < 0.01), there was no significant interaction between sampling date and farming system (PERMANOVA, uJ and BC, *p* > 0.05; Table 1). There was no significant interaction between sampling date and farming system in soil microbial community clustering (PERMANOVA, BC and uJ, *p* = 1 BF).

Within the CNT and OG farming systems, a large number of OTUs were commonly sampled across the growing season: CNT samples shared 4,530 (53.0% of total OTUs), OG shared 4,536 (53.1% of total OTUs), and OT shared 4,207 OTUs (49.2% of total OTUs). All farming systems shared 3,555 OTUs in April (54.3% of total), 3,086 OTUs in May (47.2% of total), 2,973 OTUs in early June (45.4% of total), 3,025 in late June (46.2% of total), and 2,043 in July (23.9% of total).

Random forest classification was not able to predict bacterial communities based on farming system (Fig 4; OOB estimate of error rate: 48%) or time point (not shown; OOB estimate of error rate: 69%) with reliably high accuracy. OT plots contained more putative nitrogen-fixing bacterial genera than OG or CNT (Fig. 5A). *Arthrobacter* was more abundant in organic plots across all time points, and particularly OT plots, though CNT plots contained more *Flavobacterium* and an unclassified 97%-cutoff OTU in the Bradyrhizobiaceae family (Fig. 5A). All but the least abundant genera were significantly (*p* < 0.05) differential for farming system (Fig. 5B). Soil moisture, soil temperature, and total and individual weed species biomas were important predictors of *Arthrobacter* abundance (Fig. 5C).

**Figure 4.**
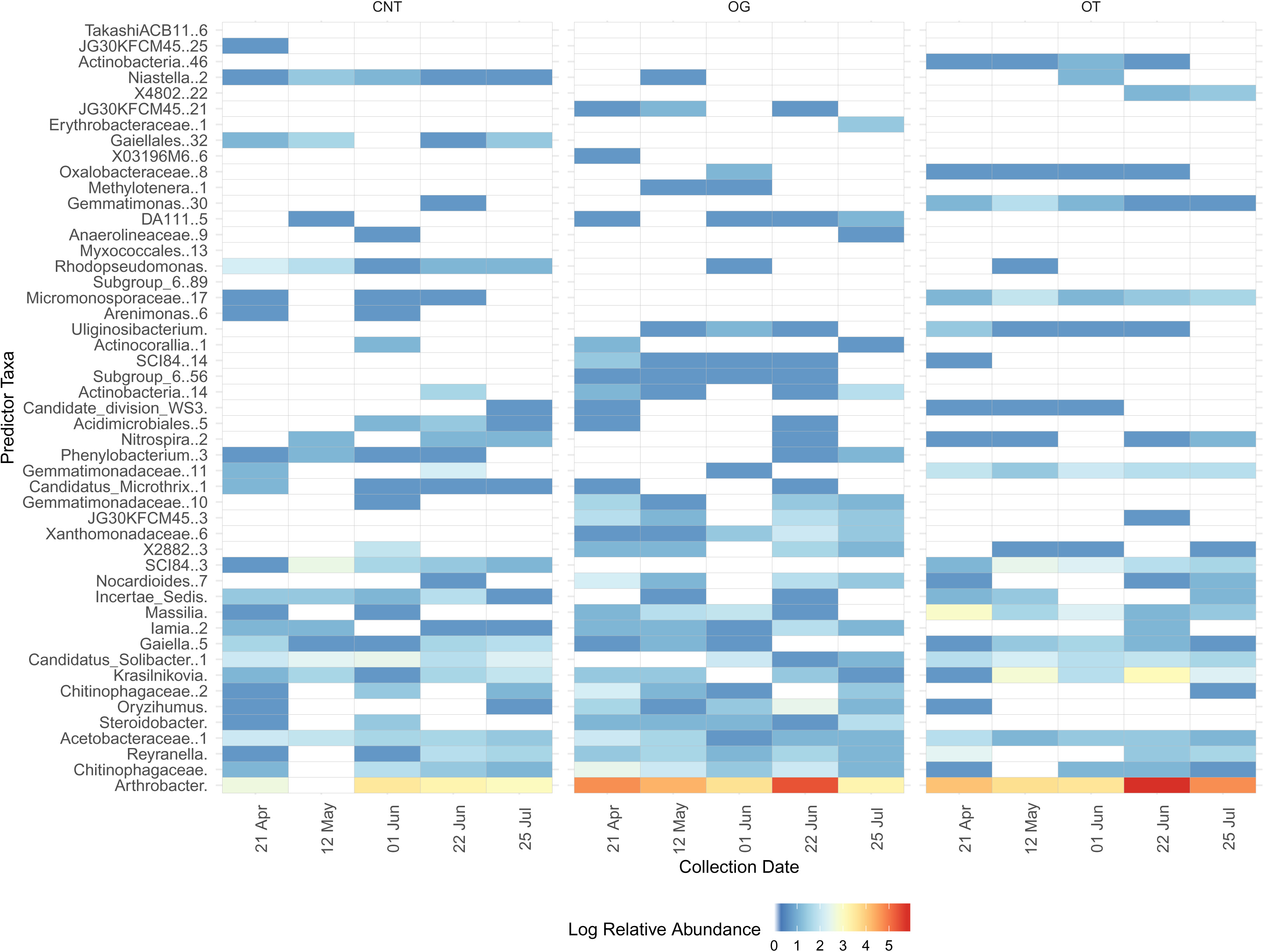
Relative abundance of 97% cutoff OTUs discriminatory to farming system [conventional (CNT), organic grazed (OG), and organic tilled (OT)], over the 2016 growing season. Samples are sorted along the x-axis by farming system and then by time.

**Figure 5.**
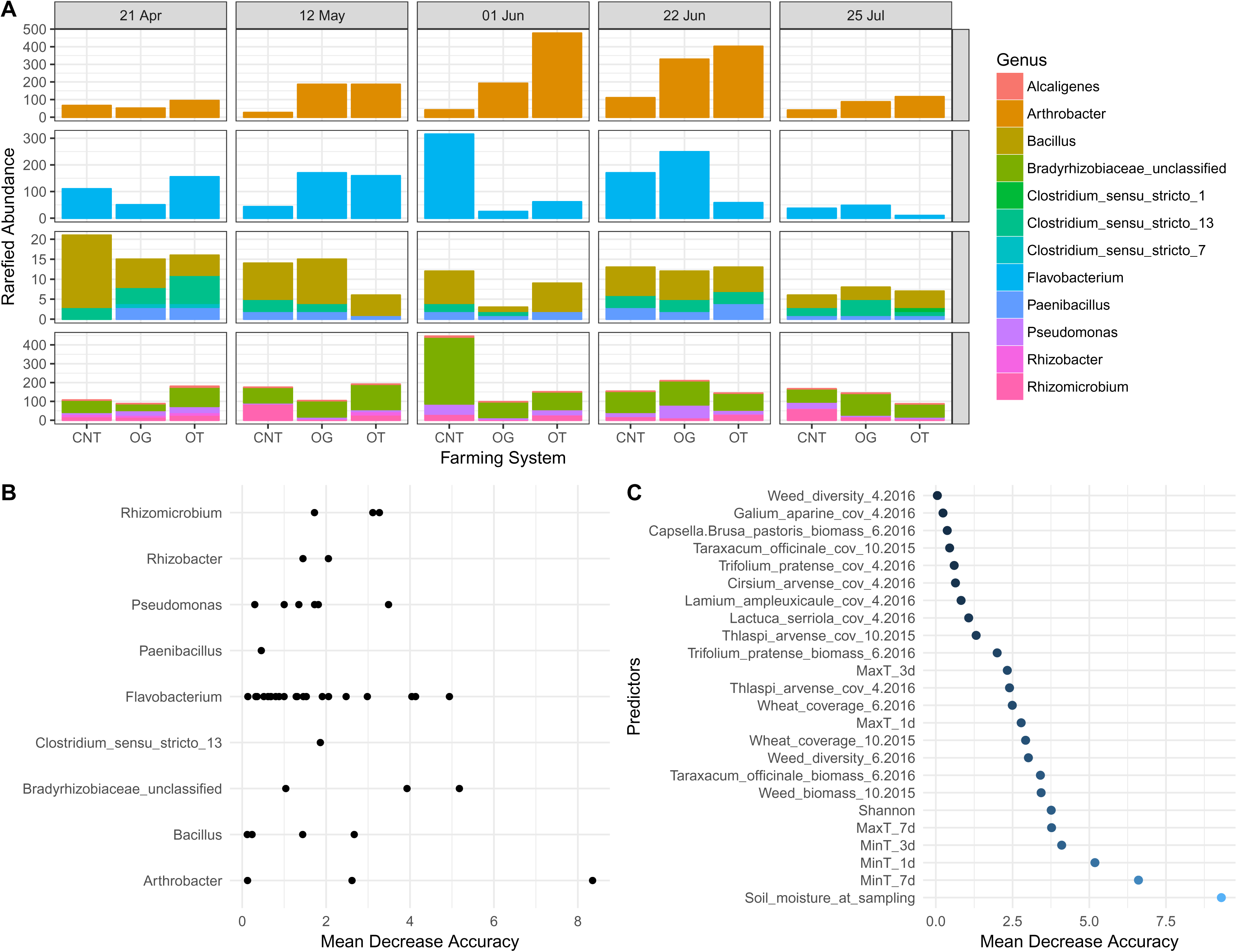
(A) Rarefied abundance of putative nitrogen-fixing bacterial genera in soil from conventional (CNT), organic grazed (OG), and organic tilled (OT) farming systems, over the 2016 growing season. (B) Importance of putative-nitrogen fixing species in discriminating between farming system. (C) Importance of factors in explaining abundance of *Arthrobacter* species in agricultural soil, including biomass or plot coverage (cov) of plants at different times, as well as minimum or maximim soil temperature (T).

### 3.2 Plant community and environmental variables

Biotic and abiotic variables affecting the soil bacterial communities included farming system, sampling date, minimum and maximum temperature on the day of sampling, total weed diversity on both April and June 2016, *Taraxacum officinale* percent coverage the previous fall in October 2015, *Lamium ampleuxicaule* percent coverage in April 2016, and *T. arvense* percent coverage in April 2016 (ANOVA, *p* < 0.05; Fig. 6).

**Figure 6.**
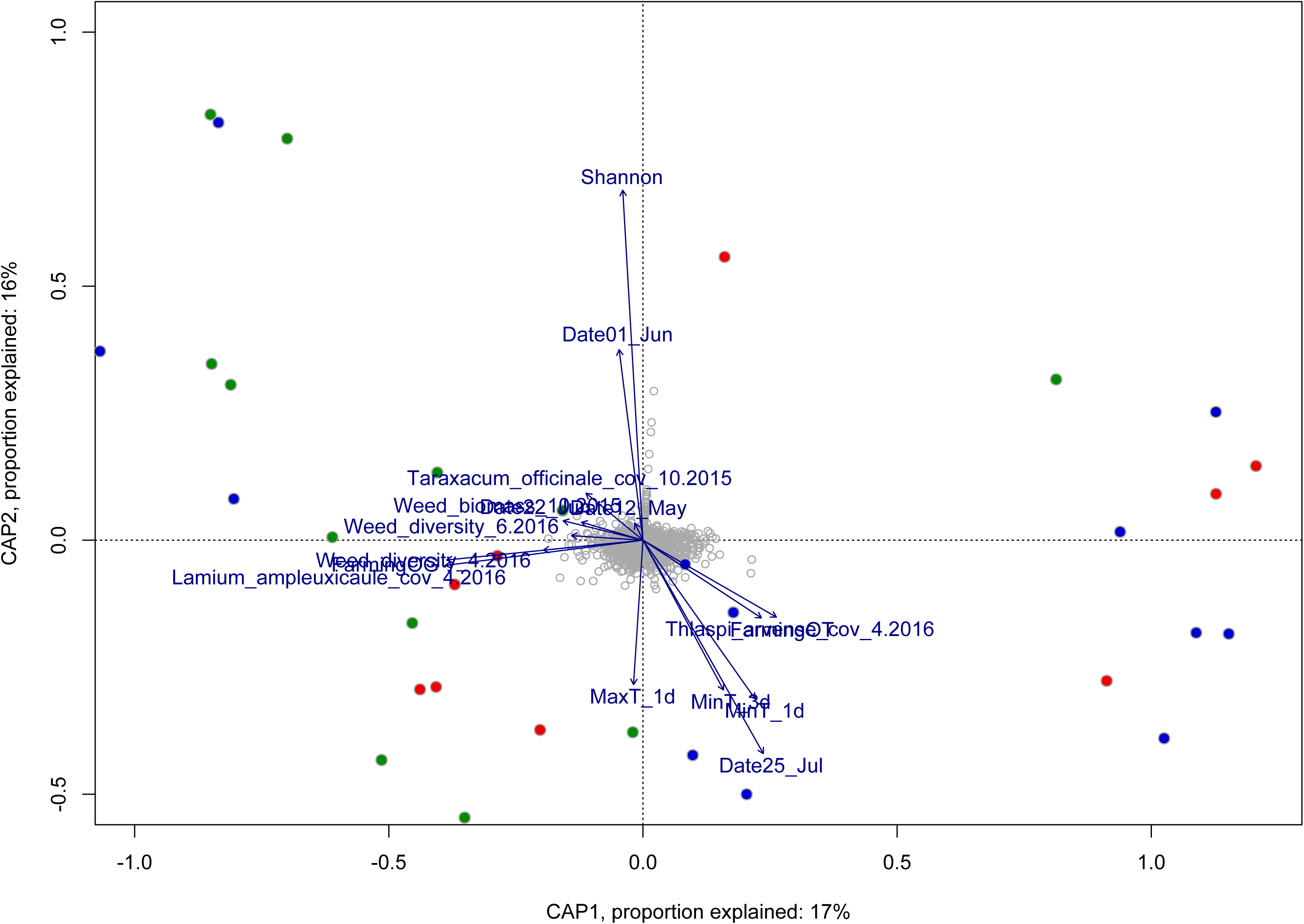
Distance-based redundancy analysis (dbRDA) of significant factors determining Hellinger-transformed soil microbial community data from conventional (CNT, red), organic grazed (OG, green), and organic tilled (OT, blue) systems. The model was significant: ANOVA, F = 1.4191, *p* = 0.001, as were the axes: CAP1, F = 6.9957, *p* = 0.002; CAP2, F = 6.5696, *p* = 0.002. Significant factors (ANOVA, *p* < 0.05) are listed in the main text.

Only weak (−0.4 to 0.4) Pearson’s correlations existed between bacterial Shannon diversity and various bacterial and weed species (Fig. S3, *p* < 0.05), likely because most samples maintained a high Shannon Index (range 5.2 - 6.6). When separated out by sampling date, the strength of the correlations between the top soil bacterial OTUs in early June (Fig. S4) and late June (Fig. S5, *p* < 0.05) increased. Total Shannon diversity and abundance of many of the most-abundant bacterial OTUs were negatively correlated with increased soil temperature at respective date of sampling (Fig. S6, *p* < 0.05). However, *Arthrobacter*, *Skermanella*, *Sphingomonas*, Comamonadaceae, *Bacteroides*, *Arenimonas*, and *Microvirga* were positively correlated with soil temperature (Fig. S6, *p* < 0.05). Soil moisture on the day of sampling was positively correlated with the most-abundant bacterial OTUs, but showed a weakly-positive correlation with Shannon diversity (Fig. S6, *p* < 0.05).

Soil pH was negatively associated with a number of bacterial genera, as was wheat protein (Fig. S7, Pearson’s corelations, *p* < 0.05). Soil pH was lower in CNT than OG (ANOVA, p < 0.05); CNT (mean 5.9), OG (mean 7.00), OT (mean 6.77). Nitrate was negatively associated with the putative genus 480-2, an unnamed clade in the Actinobacteria phylum (Fig. S7). Organic matter was not correlated with any of the most-abundant bacterial OTUs identified in July soils, but nitate was strongly negatively associated with the putative genus 480-2 (order Solirubrobacterales, phylum Actinobacteria) (Fig. S7).

## 4 Discussion

Agricultural production in the Great Plains of North America is strongly tied to seasonality, which alters temperature, moisture, solar radiation, and plant phenology. Wheat (*Triticum aestivum* L.) is the most widely-planted crop, and understanding the dynamics between production, soil microbial communities, and farming system, separately and with respect to seasonality, is important to continued sustainability. This research assessed the impact of management systems and time within the growing season on the status of bacterial communities in three contrasting dryland farming systems: a chemically-managed no-till system and two organic farming systems that were maintained for four years under USDA-organic prescribed conditions.

### 4.1 Farming system over the growing season

Farming systems selected for different bacterial communities, which has been well-established at single time-points, however we did not find significant farming system x time effects. Transitioning a farm to a USDA organic-certified system takes just three years to complete legally, yet it has been noted that soil microbial systems may take years to decades to transition to a new, stable community (Chaudhry et al., 2012; Hartmann et al., 2015; Sayer et al., 2017; Stagnari et al., 2014). Thus, the studied soil bacterial communities can be considered to be in transition, but, to our knowledge, the changes in soil microbial communities occurring during that transition or in recently certified organic system are unknown. However, the relevance of our study resides in the fact that the sampled organic fields represent certified systems from which farmers could perceive a significant economic premium (Lawrence et al., 2018; Miller et al., 2007). Variations in the soil bacterial community, including a reduction in total taxonomic diversity and a shift in abundance towards particular taxa, resulting from a change in management system occurs on a shorter time scale, (Lupwayi et al., 2004; Stagnari et al., 2014) compared with soil fungal communities (Stagnari et al., 2014). Nevertheless neither soil bacterial and fungal communities may reach a steady state for a long time, if ever, depending on the specific disturbance regime associated with in the agricultural system where they occur. Further, even after just four years under these management practices, we saw changes in the soil bacterial community.

In this study, soil from the OG system exhibited less variation between plots in bacterial species richness and moisture later into the growing season than the plots from the OT system, and a neutral pH compared to the CNT systems. In July, more dispersion was observed in community clustering, and in particular, OT plots had highly variable mean evenness and richness between field replicates in July. This suggests a random divergence of the bacterial community rather than a selective pressure in those plots at that time. These differences may suggest that farming system may mediate the effects of the environment, per our third hypothesis. Yet, the lack of strong community divergence by farming system would suggest that the effects of environment may supersede the selective effects of farming, or else that the farming systems, though past USDA certification, had not been implemented long enough to significantly alter bacterial communities in the soil. However, many of our time x farming system comparisons lacked significiance, possibly due to the variability in OT plots, or too few replicates. The effect of time within the growing season on soil microbial community dynamics is not well studied (DeBruyn et al., 2011; Marine et al., 2015).

### 4.3 Plant community and environmental effects over the growing season

July in Montana is very hot and dry, so much so that all three farming systems exhibited similarly low moisture levels. The unimodal nature of bacterial richness and eveness in all systems coincides with peak crop growth, and the reduction is July corresponds with both the weather and crop senescence, both of which would affect microbial communities. Bacterial community structure in soil varies seasonally, reflecting seasonal changes in plant phenology, solar radiation, moisture or temperature, all strong drivers of bacterial species richness (Prevost-Boure et al., 2010; Wu et al., 2016). Bacterial activity, such as nitrogen fixation, is also strongly tied to soil moisture (Koranda et al., 2013; Orr et al., 2012). In our study, soil moisture was not a significant factor structuring the entire soil bacterial community, which may reflect the prevalence of relic DNA in soils (Carini et al., 2017). However, moisture was an important predictor of the abundance of the genus *Arthrobacter*, some members of which are nitrogen-fixers (Westerberg et al., 2000), and which are reported to thrive in adverse conditions, including low moisture and high radiation conditions (Mongodin et al., 2006; SantaCruz-Calvo et al., 2013). Futher, numerous *Arthobacter* species are adept at using various carbon sources, including pesticides (Hagedorn and Holt, 1975). This is in contrast to other nitrogen-fixers, which are susceptible to pesticide toxicity (Orr et al., 2012).

Changes in weed diversity has been found to correlate with soil bacterial diversity throughout the year (Aguilera et al., 2017; Cardinale et al., 2015; Grayston et al., 1998; Ishaq et al., 2017; Whiting et al., 2001). Contrary to previous studies, (Flohre et al., 2011; Kubota et al., 2015) though, bacterial richness was positively correlated with total weed diversity only in April and weakly negatively correlated with total weed diversity in June, which was driven by the interaction between bacteria and several plant species. In agreement with the present study, *T. arvense* has been found to be associated with low bacterial diversity in soil (Whiting et al., 2001), possibly because members of the Brassicaceae family of plants produce a number of glucosinolates and other antimicrobial compounds (Pal Vig et al., 2009), which may inhibit microbial growth in the rhizosphere. Similarly, and in accordance with (Cardinale et al., 2015), *L. serricola* was associated with low soil bacterial relative abundance.

Previous studies have shown that some weed species are better at recruiting beneficial microbiota than domesticated crops, allowing them to be more competitive (Aguilera et al., 2017; Massenssini et al., 2015; Trognitz et al., 2016). Recent discussions have examined the possibility that weed species are more dependent on plant-microbial connections, while crops only need them under non-ideal growing conditions (Trognitz et al., 2016). Also, it has been shown that plants will reprioritize microbial interactions under stressful conditions (Fuchslueger et al., 2014). However, seasonal changes in moisture or nutrient availability, as well as plant phenology, dictate the need for plants to form symbiotic relationships with soil microbiota (Fuchslueger et al., 2014; Kumar et al., 2018), thus the interaction between plant species and microbial species may be uncoupled at certain points of the year, i.e. at plant senesence.

### 4.4 Summary

In the context of these receent studies, our results raise speculative but interesting questions for the sustainability of managed systems; how does soil bacterial diversity impact crop-weed competitive interactions under stressed and ideal environmental situations? Does the degree of these dependeces vary across farming systems (Johnson et al., 2017)? What are the mechanisms driving the impact of soil microbial communities on plant growth and multi-trophic interactions? Local environmental conditions strongly affect any practical applications aimed at improving soil diversity and functionality, especially in semi-arid regions where abiotic stress and seasonal variability in temperature and water availability drive primary production. Thus, it is imperative to incorporate seasonality into studies on the microbial ecology of agricultural systems.

## 5 Acknowledgements

The authors would like to thank Kyla Crisp, Madison Nixon, Tessa Scott, Rachel Flowers, Ali Thornton, and Lazaro Vinola for their assistance maintaining the plots and collecting samples; Devon Ragen for sheep maintenance; Drs. Pat Hatfield and Perry Miller for farm administration; and Sarah Olivo for DNA sequencing. This work was supported by the USDA NIFA Organic Transitions (ORG) program (Grant MONB00128).

