## Supplemental Tables and Figures for "Rhizosphere bacterial communities of wheat vary across the growing season and among dryland farming systems"

### Supplementary Tables

**Table S1 Temperature and precipitation in Bozeman, MT during the 2016 growing season and 1981 – 2010 growing seasons.**

| | April | May | June | July |
|---|---|---|---|---|
| 2016 Max °C | 25.5 | 28.3 | 35 | 36.7 |
| 2016 Min °C | 6.1 | 4.4 | 16.1 | 18.9 |
| 1981 - 2010 Max °C | 13.6 | 18.6 | 22.7 | 28.2 |
| 1981 - 2010 Min °C | -1.1 | 3.3 | 6.7 | 9.8 |
| 2016 Mean Precipitation (mm) [1] | 53 | 53 | 55 | 22 |
| 1981 - 2010 Mean Precipitation (mm) [2] | 45 | 71 | 71 | 36 |

[1] Data from Montana.gov Station Reports, 2016

[2] Data from U.S. Climate Data, PRISM Climate Group

**Table S2 Soil properties measured at wheat harvest, July 25, 2016.**

| Sample | Farming | pH | Organic matter | Nitrate (ppm) | P, Olsen (ppm) | K (ppm) |
|---|---|---|---|---|---|---|
| CC_725_n_a_1062 | CNT | 6.4 | 5.3 | 2.5 | 51 | 510 |
| CC_725_n_a_3052 | CNT | 5.4 | 5.7 | 4.5 | 37 | 335 |
| OG_725_n_a_1022 | OG | 7.3 | 6 | 10 | 53 | 851 |
| OG_725_n_a_2092 | OG | 7.1 | 5.8 | 1.5 | 44 | 350 |
| OG_725_n_a_3152 | OG | 6.6 | 6.1 | 2 | 45 | 389 |
| OT_725_n_a_1142 | OT | 7.9 | 6.1 | 3 | 38 | 263 |
| OT_725_n_a_2022 | OT | 6.6 | 7.3 | 7 | 90 | 813 |
| OT_725_n_a_3102 | OT | 5.8 | 6.3 | 0.5 | 37 | 466 |

**Table S3 Comparison values for relative abundance for phyla in soil from conventional (CNT), organic grazed (OG), and organic tilled (OT) systems.** Significance was determined by Wilcoxon rank, $p < 0.05$ = *, $p \leq 0.01$ = **, $p \leq 0.001$ = ***. Relative abundance is visualized in Figure S2.

| | CNT - OG | | | | | CNT - OT | | | | | OG - OT | | | | |
|---|---|---|---|---|---|---|---|---|---|---|---|---|---|---|---|
| Phyla | 04.21.2016 | 05.12.2016 | 06.01.2016 | 06.22.2016 | 07.25.2016 | 04.21.2016 | 05.12.2016 | 06.01.2016 | 06.22.2016 | 07.25.2016 | 04.21.2016 | 05.12.2016 | 06.01.2016 | 06.22.2016 | 07.25.2016 |
| Acidobacteria | - | - | - | - | - | - | - | * | - | - | - | - | - | - | - |
| Actinobacteria | - | - | - | - | - | - | * | * | - | - | - | - | - | - | * |
| Armatimonadetes | - | - | - | * | - | - | - | - | - | - | - | - | - | - | - |
| BD1-5 | - | - | * | - | - | - | - | - | - | - | - | - | * | - | - |
| Candidate BRC1 | - | - | - | - | - | - | - | - | - | - | - | - | - | ** | - |
| Candidate TM7 | - | - | - | - | - | - | - | - | - | * | - | - | - | - | - |
| Candidate WS3 | - | - | - | - | - | - | - | * | - | - | - | - | - | - | - |
| Chlorobi | - | - | - | - | - | - | - | - | - | - | - | - | * | - | - |
| Chloroflexi | ** | - | - | - | - | - | - | - | - | - | - | * | - | - | - |
| Cyanobacteria | - | - | * | - | * | - | - | - | - | - | * | - | - | - | - |
| Deinococcus-Thermus | - | * | - | - | - | ** | - | - | - | - | - | - | - | - | - |
| Fibrobacteres | * | * | - | - | - | - | - | - | - | - | - | - | - | - | * |
| Firmicutes | - | * | * | - | - | - | - | - | - | - | - | - | - | - | - |
| Proteobacteria | - | - | - | - | - | - | - | * | - | - | - | *** | - | - | - |
| SHA-109 | - | - | - | - | * | * | - | - | - | - | * | - | - | - | - |
| SM2F11 | * | - | - | - | - | * | - | - | - | - | - | - | - | - | - |
| Verrucomicrobia | * | - | - | - | - | - | - | - | - | - | - | - | - | - | - |
| WCHB1-60 | - | - | - | * | - | - | - | - | - | - | - | - | - | ** | - |

Euryarchaeota (Archaea), Bacteroidetes, Caldiserica, Chlamydiae, Elusimicrobia, Fusobacteria, Gemmatimonadetes, Lentisphaerae, Nitrospirae, Planctomycetes, Spirochaetae, Tenericutes, Thermotogae, and Candidate phyla GOUTA4, JL-ETNP-Z39, OD1, OP11, OP3, SR1, TM6, WS6, and WD272, were not significantly different between any system at any timepoint.

**Table S4 Pairwise PERMANOVA model output of the effects of farming systems on soil microbial communities for OTUs at a 97% genetic cutoff. P-values were Bonferroni corrected.**

| Bray-Curtis | | | Unweighted Jaccard | | |
|---|---|---|---|---|---|
| | CNT | OG | | CNT | OG |
| OG | 0.035 | - | OG | 0.034 | - |
| OT | 0.14 | 0.14 | OT | 0.160 | 0.160 |

**Table S5 Pairwise PERMANOVA model output of the effects of date of sampling s on soil microbial communities for OTUs at a 97% genetic cutoff. P-values were Bonferroni corrected.**

| Bray-Curtis | | | | | Unweighted Jaccard | | | | |
|---|---|---|---|---|---|---|---|---|---|
| | 01 Jun | 12 May | 21 Apr | 22 Jun | | 01 Jun | 12 May | 21 Apr | 22 Jun |
| 12 May | 1 | - | - | - | 12 May | 1 | - | - | - |
| 21 Apr | 0.645 | 1 | - | - | 21 Apr | 1 | 1 | - | - |
| 22 Jun | 0.167 | 1 | 1 | - | 22 Jun | 1 | 1 | 1 | - |
| 25 Jul | 0.014 | 0.37 | 0.295 | 0.022 | 25 Jul | 0.024 | 1 | 1 | 0.227 |

### Supplementary Figures

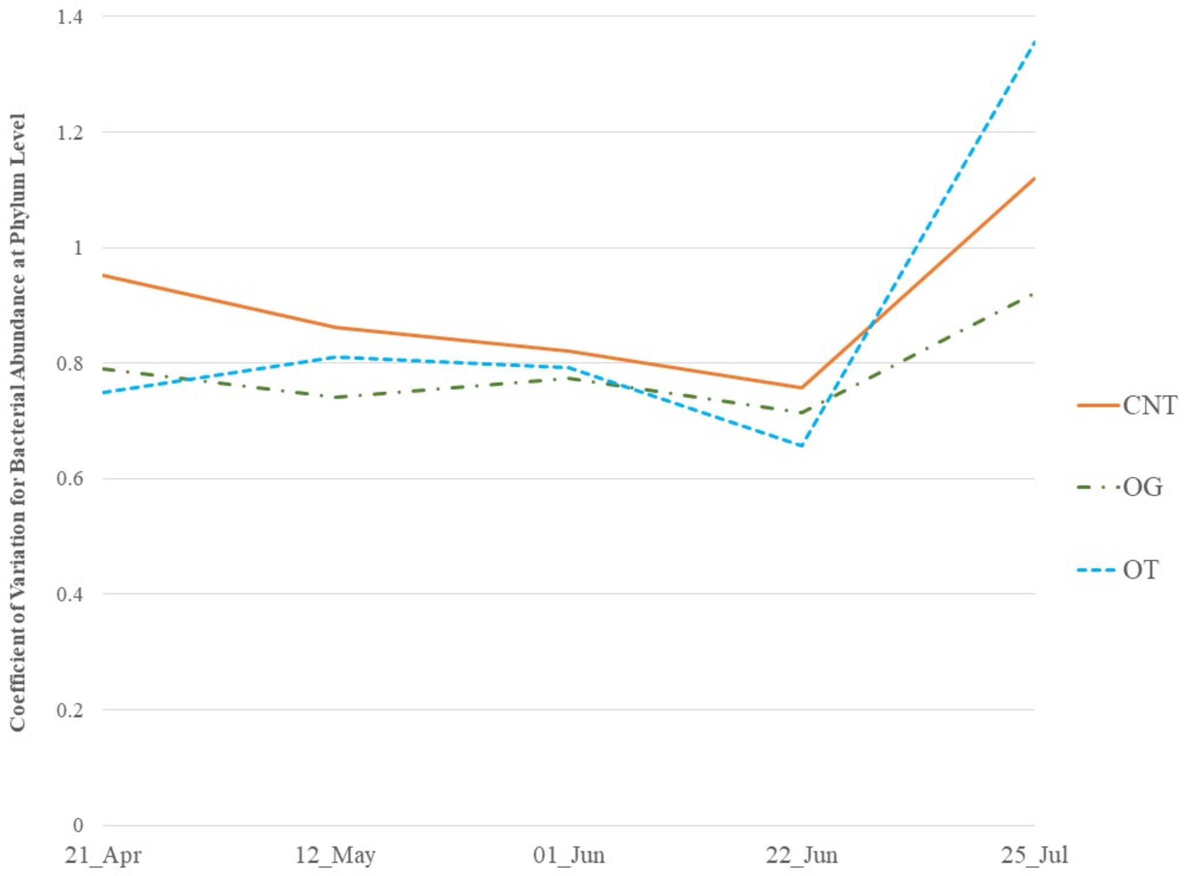

**Fig S1 Mean coefficient of variation of phylum-level relative abundance for bacterial communities in soil from conventional (CNT), organic grazed (OG), and organic tilled (OT) systems), over the 2016 growing season.**

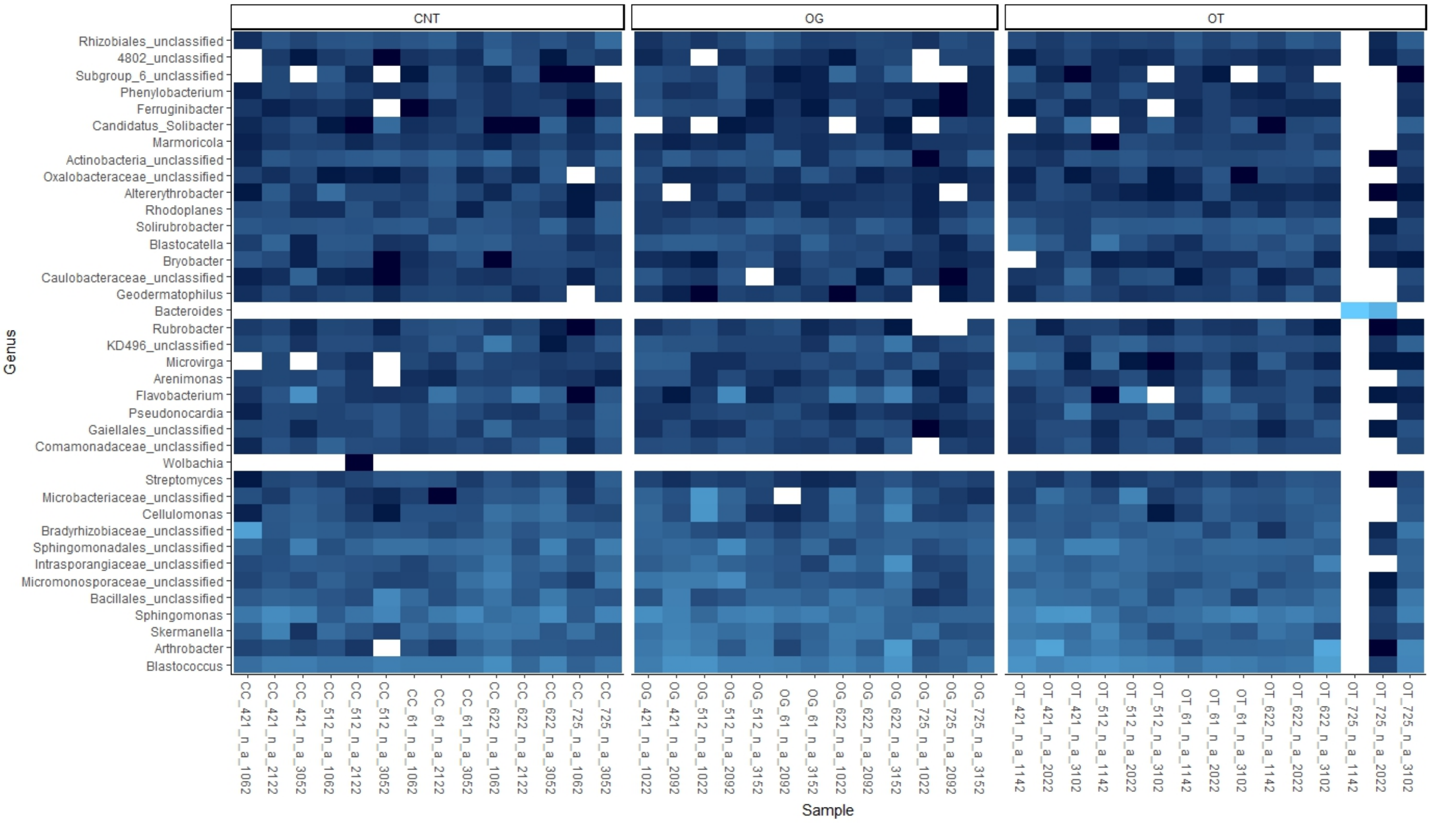

CNT
OG
OT
Genus
Rhizobiales_unclassified
4802_unclassified
Subgroup_6_unclassified
Phenylobacterium
Ferruginibacter
Candidatus_Solibacter
Marmoricola
Actinobacteria_unclassified
Oxalobacteraceae_unclassified
Altererythrobacter
Rhodoplanes
Solirubrobacter
Blastocatella
Bryobacter
Caulobacteraceae_unclassified
Geodermatophilus
Bacteroides
Rubrobacter
KD496_unclassified
Microvirga
Arenimonas
Flavobacterium
Pseudonocardia
Gaiellales_unclassified
Comamonadaceae_unclassified
Wolbachia
Streptomyces
Microbacteriaceae_unclassified
Cellulomonas
Bradyrhizobiaceae_unclassified
Sphingomonadales_unclassified
Intrasporangiaceae_unclassified
Micromonosporaceae_unclassified
Bacillales_unclassified
Sphingomonas
Skermanella
Arthrobacter
Blastococcus
CC_421_n_a_1062
CC_421_n_a_2122
CC_421_n_a_3052
CC_512_n_a_1062
CC_512_n_a_2122
CC_512_n_a_3052
CC_61_n_a_1062
CC_61_n_a_2122
CC_61_n_a_3052
CC_622_n_a_1062
CC_622_n_a_2122
CC_622_n_a_3052
CC_725_n_a_1062
CC_725_n_a_3052
OG_421_n_a_1022
OG_421_n_a_2092
OG_512_n_a_1022
OG_512_n_a_2092
OG_512_n_a_3152
OG_61_n_a_2092
OG_61_n_a_3152
OG_622_n_a_1022
OG_622_n_a_2092
OG_622_n_a_3152
OG_725_n_a_1022
OG_725_n_a_2092
OG_725_n_a_3152
OT_421_n_a_1142
OT_421_n_a_2022
OT_421_n_a_3102
OT_512_n_a_1142
OT_512_n_a_2022
OT_512_n_a_3102
OT_61_n_a_1142
OT_61_n_a_2022
OT_61_n_a_3102
OT_622_n_a_1142
OT_622_n_a_2022
OT_622_n_a_3102
OT_725_n_a_1142
OT_725_n_a_2022
OT_725_n_a_3102
Sample

**Figure S2 Relative abundance of species-level bacterial OTUs in soil from conventional (CNT), organic grazed (OG), and organic tilled (OT) systems), over the 2016 growing season.** Samples are sorted along the x-axis by farming system and then by time.

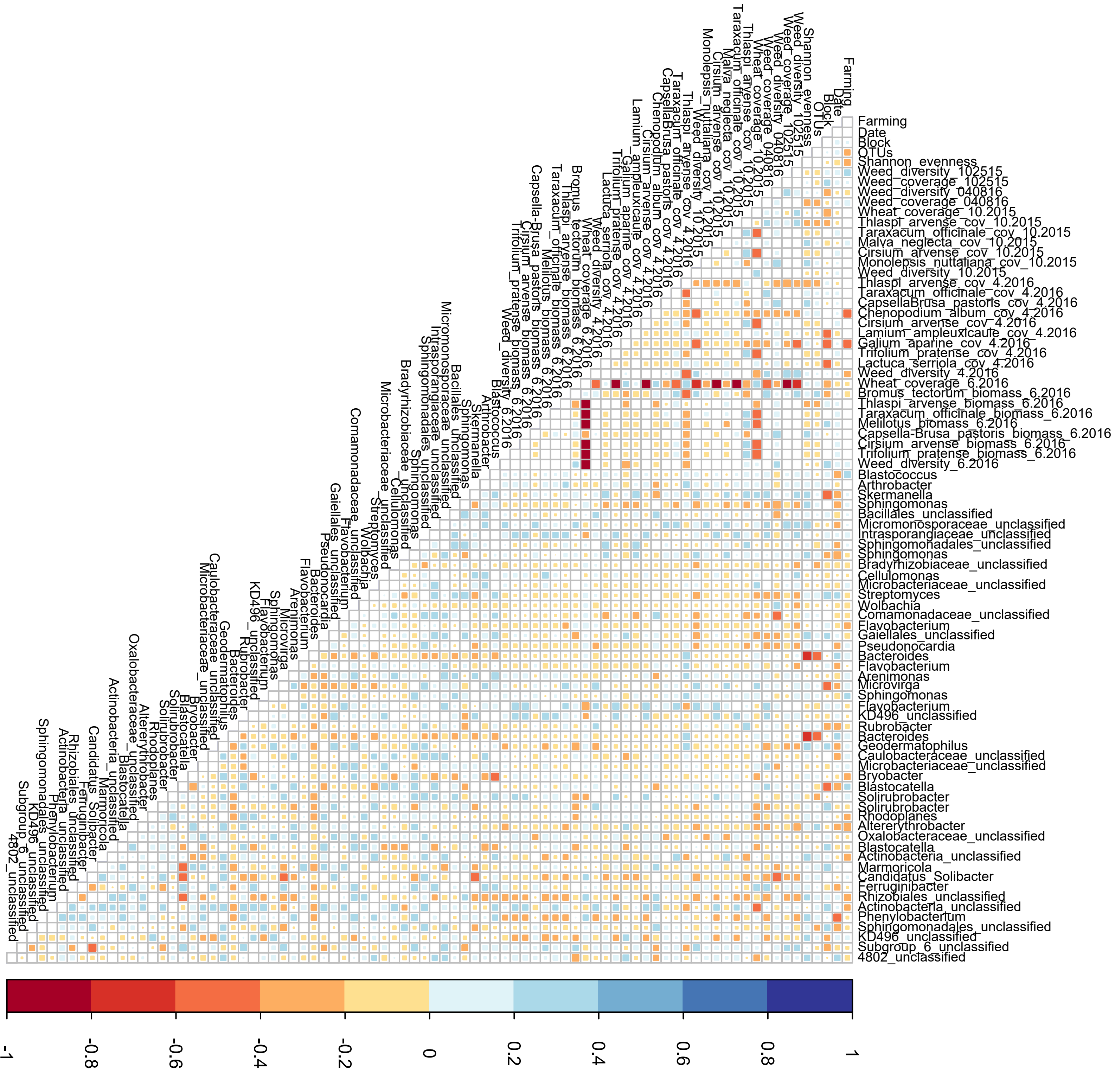
Farming
Date
Block
OTUs
Shannon_evenness
Weed_diversity_102515
Weed_coverage_102515
Weed_diversity_040816
Weed_coverage_040816
Wheat_coverage_10.2015
Thlaspi_arvense_cov_10.2015
Taraxacum_officinale_cov_10.2015
Malva_neglecta_cov_10.2015
Cirsium_arvense_cov_10.2015
Monolepsis_nuttaliana_cov_10.2015
Weed_diversity_10.2015
Thlaspi_arvense_cov_4.2016
Taraxacum_officinale_cov_4.2016
CapsellaBrusa_pastoris_cov_4.2016
Chenopodium_album_cov_4.2016
Cirsium_arvense_cov_4.2016
Lamium_ampleuxicaule_cov_4.2016
Galium_aparine_cov_4.2016
Trifolium_pratense_cov_4.2016
Lactuca_serriola_cov_4.2016
Weed_diversity_4.2016
Wheat_coverage_6.2016
Bromus_tectorum_biomass_6.2016
Thlaspi_arvense_biomass_6.2016
Taraxacum_officinale_biomass_6.2016
Melilotus_biomass_6.2016
Capsella-Brusa_pastoris_biomass_6.2016
Cirsium_arvense_biomass_6.2016
Trifolium_pratense_biomass_6.2016
Weed_diversity_6.2016
Blastococcus
Arthrobacter
Skermanella
Sphingomonas
Bacillales_unclassified
Micromonosporaceae_unclassified
Intrasporangiaceae_unclassified
Sphingomonadales_unclassified
Sphingomonas
Bradyrhizobiaceae_unclassified
Cellulomonas
Microbacteriaceae_unclassified
Streptomyces
Wolbachia
Comamonadaceae_unclassified
Flavobacterium
Gaiellales_unclassified
Pseudonocardia
Bacteroides
Flavobacterium
Arenimonas
Microvirga
Sphingomonas
Flavobacterium
KD496_unclassified
Rubrobacter
Bacteroides
Geodermatophilus
Caulobacteraceae_unclassified
Microbacteriaceae_unclassified
Bryobacter
Blastocatella
Solirubrobacter
Solirubrobacter
Rhodoplanes
Altererythrobacter
Oxalobacteraceae_unclassified
Blastocatella
Actinobacteria_unclassified
Marmoricola
Candidatus_Solibacter
Ferruginibacter
Rhizobiales_unclassified
Actinobacteria_unclassified
Phenylobacterium
Sphingomonadales_unclassified
KD496_unclassified
Subgroup_6_unclassified
4802_unclassified
-1
-0.8
-0.6
-0.4
-0.2
0
0.2
0.4
0.6
0.8
1

**Figure S3 Pearson's correlations of weed diversity data to species-level bacterial OTUs in soil from conventional (CNT), organic grazed (OG), and organic tilled (OT) systems, over the 2016 growing season.** Significance was determined by Wilcoxon rank, $p < 0.05$, and only significant values are shown.

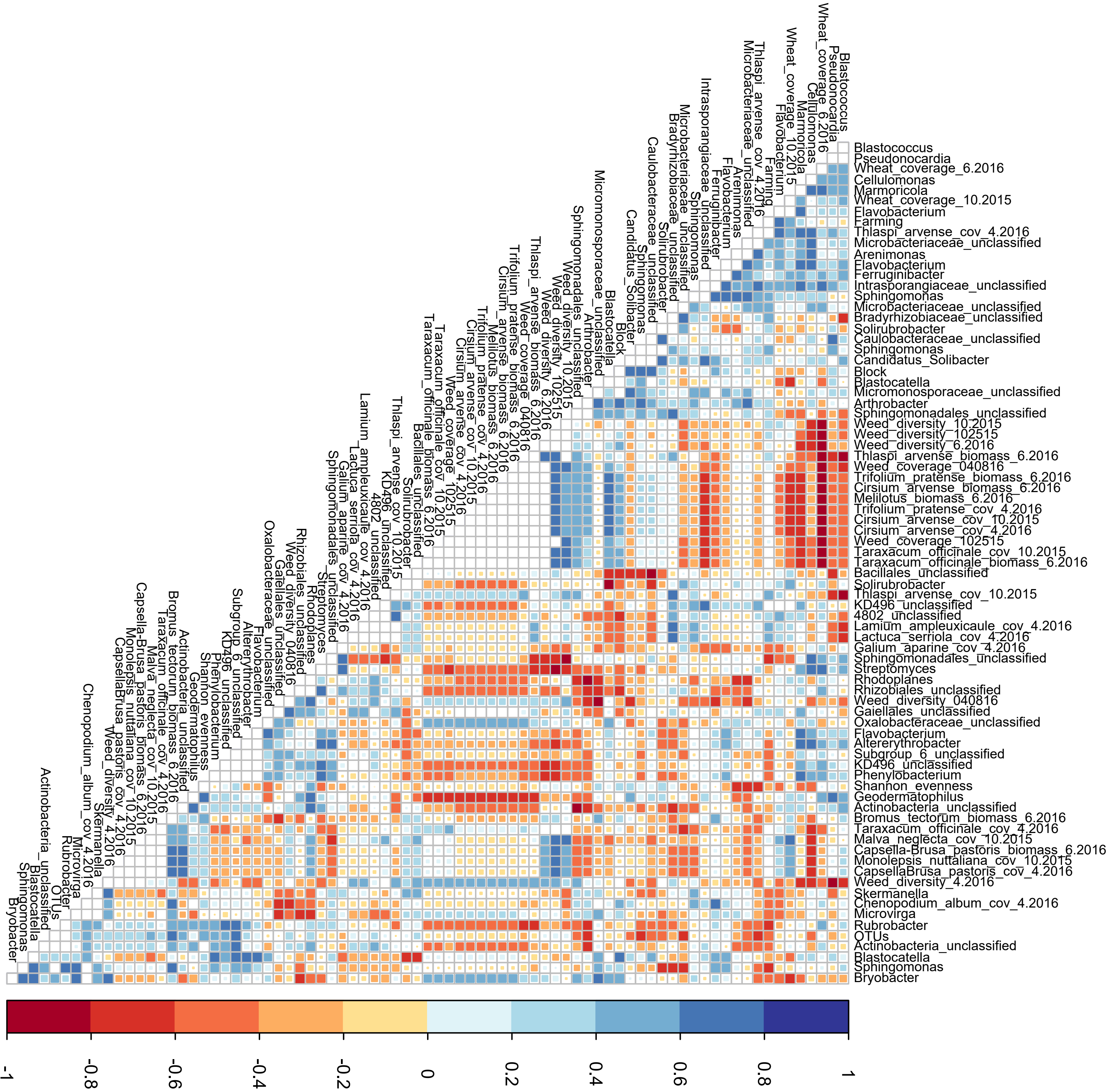
Blastococcus
Pseudonocardia
Wheat_coverage_6.2016
Cellulomonas
Marmoricola
Wheat_coverage_10.2015
Flavobacterium
Farming
Thlaspi_arvense_cov_4.2016
Microbacteriaceae_unclassified
Arenimonas
Flavobacterium
Ferruginibacter
Intrasporangiaceae_unclassified
Sphingomonas
Microbacteriaceae_unclassified
Bradyrhizobiaceae_unclassified
Solirubrobacter
Caulobacteraceae_unclassified
Sphingomonas
Candidatus_Solibacter
Block
Blastocatella
Micromonosporaceae_unclassified
Arthrobacter
Sphingomonadales_unclassified
Weed_diversity_10.2015
Weed_diversity_102515
Weed_diversity_6.2016
Thlaspi_arvense_biomass_6.2016
Weed_coverage_040816
Trifolium_pratense_biomass_6.2016
Cirsium_arvense_biomass_6.2016
Melilotus_biomass_6.2016
Trifolium_pratense_cov_4.2016
Cirsium_arvense_cov_10.2015
Cirsium_arvense_cov_4.2016
Weed_coverage_102515
Taraxacum_officinale_cov_10.2015
Taraxacum_officinale_biomass_6.2016
Bacillales_unclassified
Solirubrobacter
Thlaspi_arvense_cov_10.2015
KD496_unclassified
4802_unclassified
Lamium_ampleuxicaule_cov_4.2016
Lactuca_serriola_cov_4.2016
Galium_aparine_cov_4.2016
Sphingomonadales_unclassified
Streptomyces
Rhodoplanes
Rhizobiales_unclassified
Weed_diversity_040816
Gaiellales_unclassified
Oxalobacteraceae_unclassified
Flavobacterium
Altererythrobacter
Subgroup_6_unclassified
KD496_unclassified
Phenylobacterium
Shannon_evenness
Geodermatophilus
Actinobacteria_unclassified
Bromus_tectorum_biomass_6.2016
Taraxacum_officinale_cov_4.2016
Malva_neglecta_cov_10.2015
Capsella-Brusa_pastoris_biomass_6.2016
Monolepsis_nuttaliana_cov_10.2015
CapsellaBrusa_pastoris_cov_4.2016
Weed_diversity_4.2016
Skermanella
Chenopodium_album_cov_4.2016
Microvirga
Rubrobacter
OTUs
Actinobacteria_unclassified
Blastocatella
Sphingomonas
Bryobacter
-1
-0.8
-0.6
-0.4
-0.2
0
0.2
0.4
0.6
0.8
1

**Figure S4 Pearson's correlations of weed diversity data to species-level bacterial OTUs in soil in early June 2016 from conventional (CNT), organic grazed (OG), and organic tilled (OT) systems.** Significance was determined by Wilcoxon rank, $p < 0.05$, and only significant values are shown.

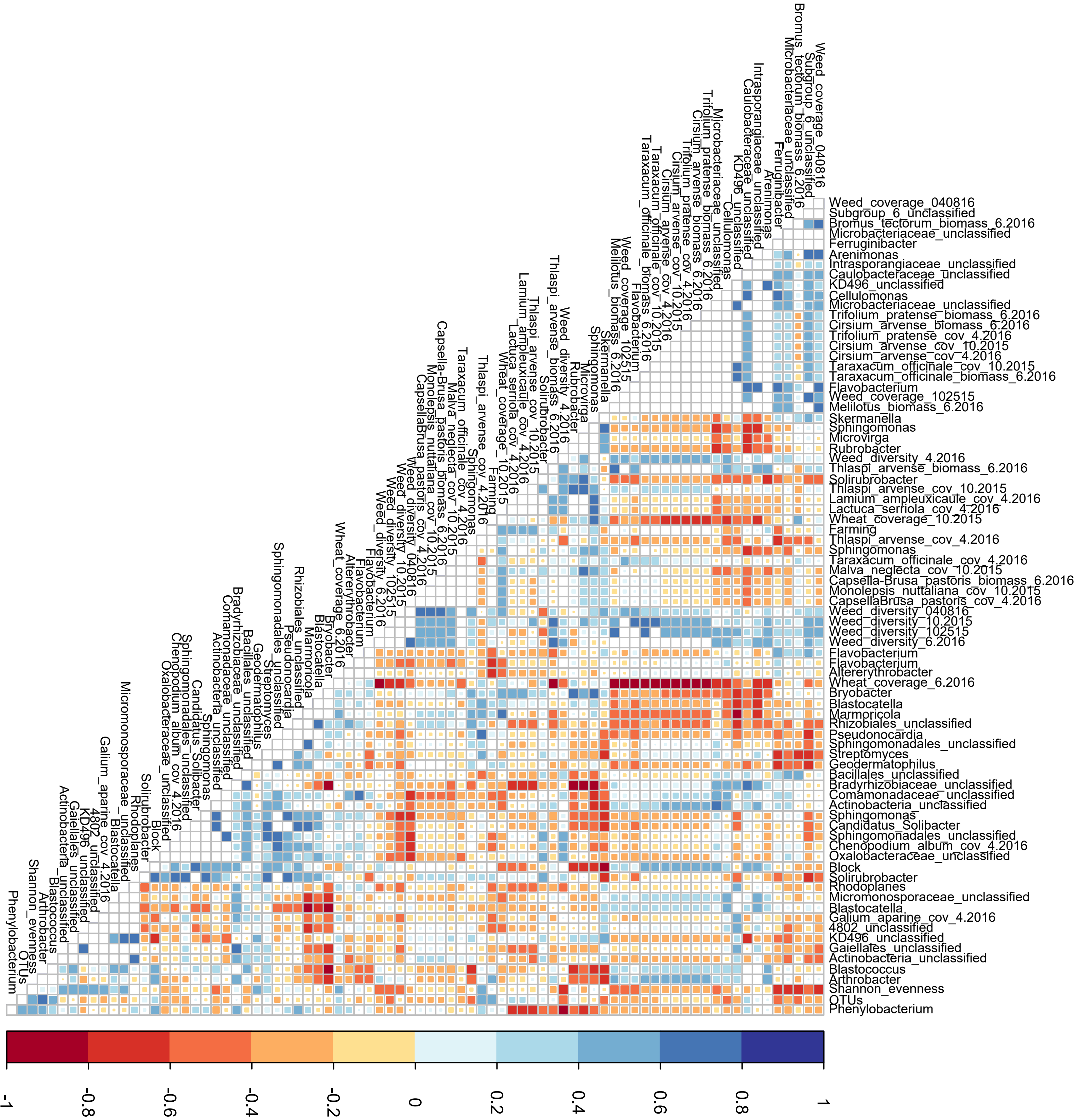
Weed_coverage_040816
Subgroup_6_unclassified
Bromus_tectorum_biomass_6.2016
Microbacteriaceae_unclassified
Ferruginibacter
Arenimonas
Intrasporangiaceae_unclassified
Caulobacteraceae_unclassified
KD496_unclassified
Cellulomonas
Microbacteriaceae_unclassified
Trifolium_pratense_biomass_6.2016
Cirsium_arvense_biomass_6.2016
Trifolium_pratense_cov_4.2016
Cirsium_arvense_cov_10.2015
Cirsium_arvense_cov_4.2016
Taraxacum_officinale_cov_10.2015
Taraxacum_officinale_biomass_6.2016
Flavobacterium
Weed_coverage_102515
Melilotus_biomass_6.2016
Skermanella
Sphingomonas
Microvirga
Rubrobacter
Weed_diversity_4.2016
Thlaspi_arvense_biomass_6.2016
Solirubrobacter
Thlaspi_arvense_cov_10.2015
Lamium_ampleuxicaule_cov_4.2016
Lactuca_serriola_cov_4.2016
Wheat_coverage_10.2015
Farming
Thlaspi_arvense_cov_4.2016
Sphingomonas
Taraxacum_officinale_cov_4.2016
Malva_neglecta_cov_10.2015
Capsella-Brusa_pastoris_biomass_6.2016
Monolepsis_nuttaliana_cov_10.2015
CapsellaBrusa_pastoris_cov_4.2016
Weed_diversity_040816
Weed_diversity_10.2015
Weed_diversity_102515
Weed_diversity_6.2016
Flavobacterium
Flavobacterium
Altererythrobacter
Wheat_coverage_6.2016
Bryobacter
Blastocatella
Marmoricola
Rhizobiales_unclassified
Pseudonocardia
Sphingomonadales_unclassified
Streptomyces
Geodermatophilus
Bacillales_unclassified
Bradyrhizobiaceae_unclassified
Comamonadaceae_unclassified
Actinobacteria_unclassified
Sphingomonas
Candidatus_Solibacter
Sphingomonadales_unclassified
Chenopodium_album_cov_4.2016
Oxalobacteraceae_unclassified
Block
Solirubrobacter
Rhodoplanes
Micromonosporaceae_unclassified
Blastocatella
Galium_aparine_cov_4.2016
4802_unclassified
KD496_unclassified
Gaiellales_unclassified
Actinobacteria_unclassified
Blastococcus
Arthrobacter
Shannon_evenness
OTUs
Phenylobacterium
-1
-0.8
-0.6
-0.4
-0.2
0
0.2
0.4
0.6
0.8
1

**Figure S5 Pearson's correlations of weed diversity data to species-level bacterial OTUs in soil in late June 2016 from conventional (CNT), organic grazed (OG), and organic tilled (OT) systems.** Significance was determined by Wilcoxon rank, $p < 0.05$, and only significant values are shown.

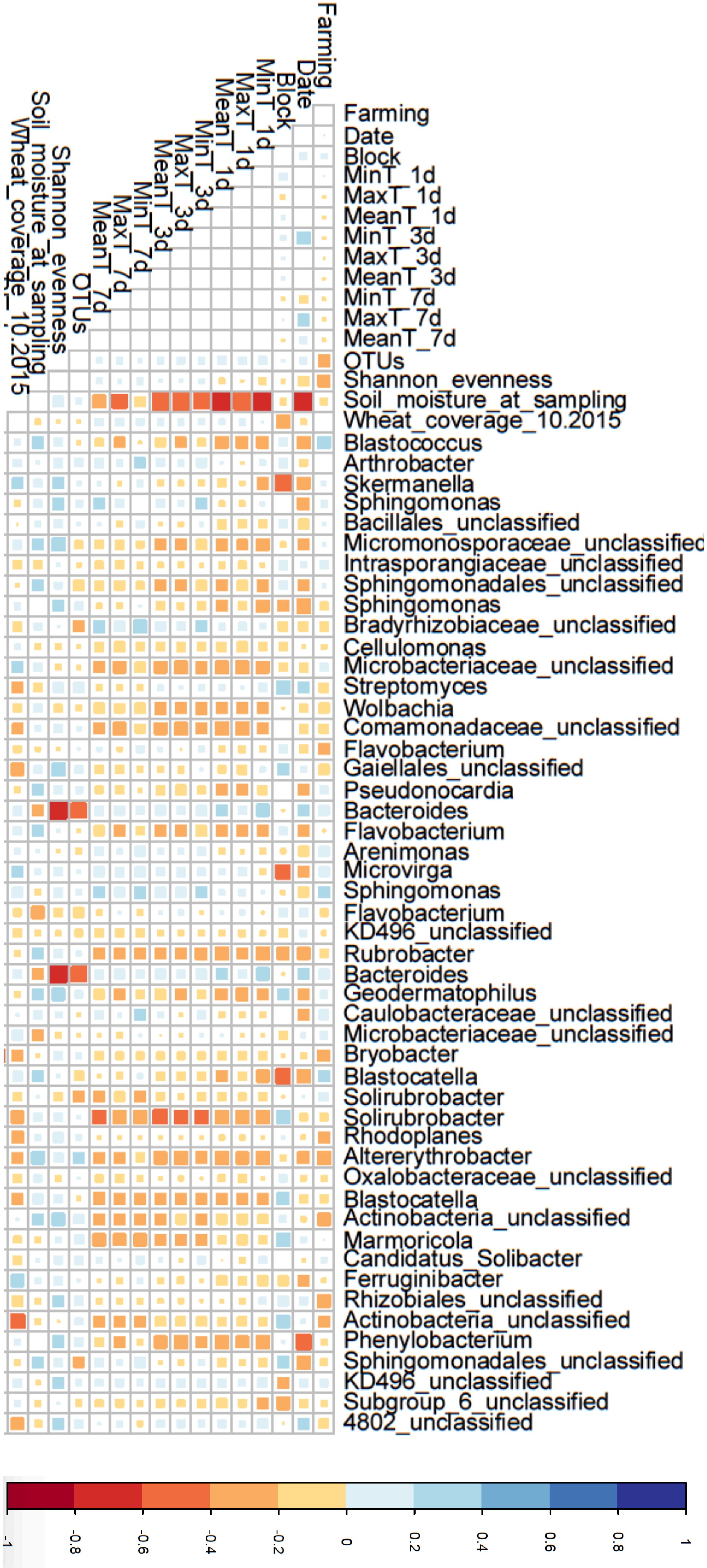
Farming
Date
Block
MinT_1d
MaxT_1d
MeanT_1d
MinT_3d
MaxT_3d
MeanT_3d
MinT_7d
MaxT_7d
MeanT_7d
OTUs
Shannon_evenness
Soil_moisture_at_sampling
Wheat_coverage_10.2015
Blastococcus
Arthrobacter
Skermanella
Sphingomonas
Bacillales_unclassified
Micromonosporaceae_unclassified
Intrasporangiaceae_unclassified
Sphingomonadales_unclassified
Sphingomonas
Bradyrhizobiaceae_unclassified
Cellulomonas
Microbacteriaceae_unclassified
Streptomyces
Wolbachia
Comamonadaceae_unclassified
Flavobacterium
Gaiellales_unclassified
Pseudonocardia
Bacteroides
Flavobacterium
Arenimonas
Microvirga
Sphingomonas
Flavobacterium
KD496_unclassified
Rubrobacter
Bacteroides
Geodermatophilus
Caulobacteraceae_unclassified
Microbacteriaceae_unclassified
Bryobacter
Blastocatella
Solirubrobacter
Solirubrobacter
Rhodoplanes
Altererythrobacter
Oxalobacteraceae_unclassified
Blastocatella
Actinobacteria_unclassified
Marmoricola
Candidatus_Solibacter
Ferruginibacter
Rhizobiales_unclassified
Actinobacteria_unclassified
Phenylobacterium
Sphingomonadales_unclassified
KD496_unclassified
Subgroup_6_unclassified
4802_unclassified
-1
-0.8
-0.6
-0.4
-0.2
0
0.2
0.4
0.6
0.8
1

**Figure S6 Pearson's correlations of soil conditions to species-level bacterial OTUs in soil from conventional (CNT), organic grazed (OG), and organic tilled (OT) systems across the 2016 growing season.** Significance was determined by Wilcoxon rank, $p < 0.05$, and only significant values are shown.

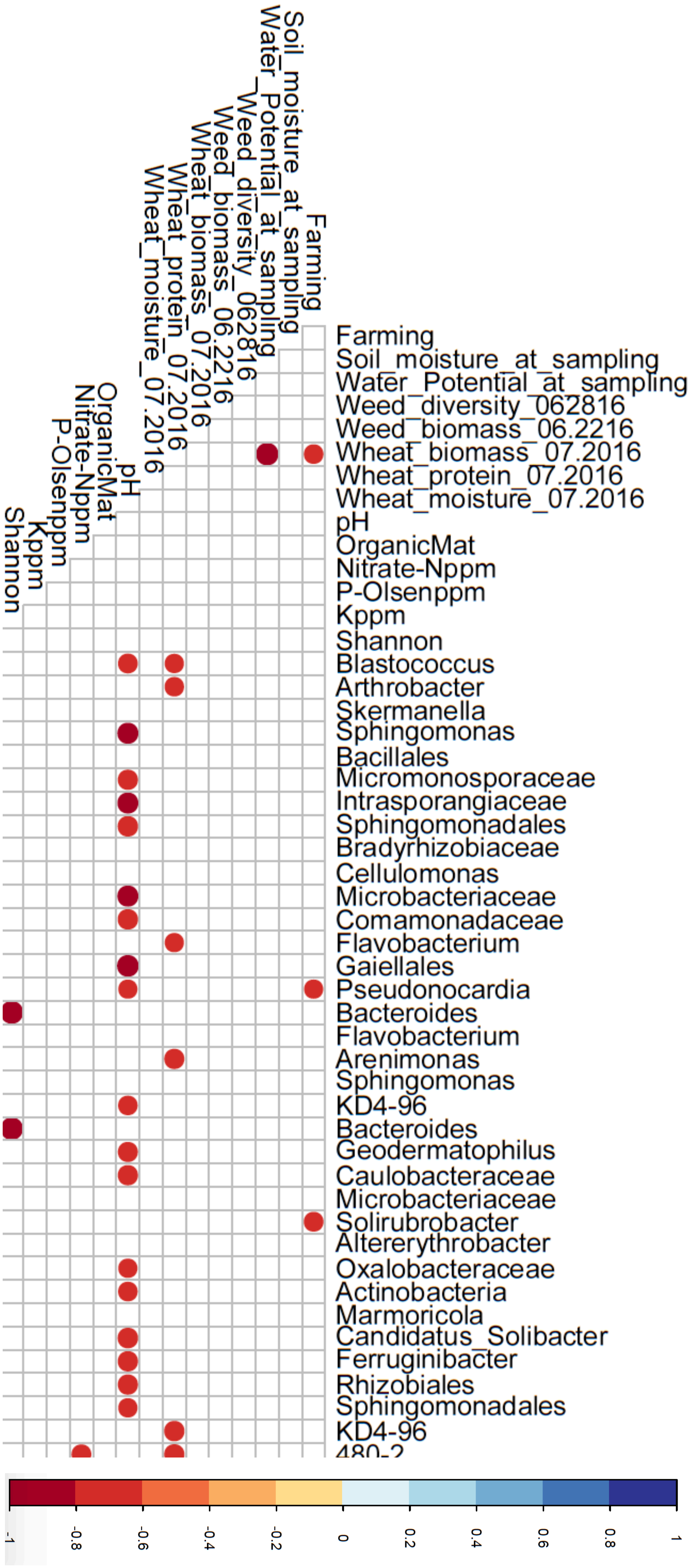

Farming
Soil_moisture_at_sampling
Water_Potential_at_sampling
Weed_diversity_062816
Weed_biomass_06.2216
Wheat_biomass_07.2016
Wheat_protein_07.2016
Wheat_moisture_07.2016
pH
OrganicMat
Nitrate-Nppm
P-Olsenppm
Kppm
Shannon
Blastococcus
Arthrobacter
Skermanella
Sphingomonas
Bacillales
Micromonosporaceae
Intrasporangiaceae
Sphingomonadales
Bradyrhizobiaceae
Cellulomonas
Microbacteriaceae
Comamonadaceae
Flavobacterium
Gaiellales
Pseudonocardia
Bacteroides
Flavobacterium
Arenimonas
Sphingomonas
KD4-96
Bacteroides
Geodermatophilus
Caulobacteraceae
Microbacteriaceae
Solirubrobacter
Altererythrobacter
Oxalobacteraceae
Actinobacteria
Marmoricola
Candidatus_Solibacter
Ferruginibacter
Rhizobiales
Sphingomonadales
KD4-96
480-2
-1
-0.8
-0.6
-0.4
-0.2
0
0.2
0.4
0.6
0.8
1

**Figure S7 Pearson's correlations of wheat and soil parameters from July 2016 to species-level bacterial OTUs in soil from conventional (CNT), organic grazed (OG), and organic tilled (OT) systems.** Significance was determined by Wilcoxon rank, $p < 0.05$, and only significant values are shown.
